# Association of RhoGEF Ect2 with Desmoplakin Supports RhoA Activity at Intercellular Junctions: Implications for Carvajal Disease

**DOI:** 10.1101/2025.05.21.655405

**Authors:** Hoda Zarkoob, Chen Yuan Kam, Jennifer Koetsier, Avinash Jaiganesh, Erin McCarthy, David Kelsell, Farah Sheikh, Lisa M. Godsel, Kathleen J. Green

**Affiliations:** Northwestern Feinberg School of Medicine, Department of Pathology, 303 E. Chicago Ave., Chicago, IL. 60611; Blizard Institute, Faculty of Medicine and Dentistry, Queen Mary University of London, London E1 4NS; University of California-San Diego, School of Medicine, Department of Medicine, La Jolla CA 92093

## Abstract

Desmoplakin (DP) is an essential component of the desmosomal adhesion complex, tethering intermediate filaments to sites of intercellular adhesion to confer mechanical integrity to tissues. As a frequent target for mutation in cardiocutaneous syndromes that vary widely in phenotype, DP’s roles as a signaling hub are rapidly emerging. Here, we identify the RhoGEF Ect2 as a previously unappreciated binding partner of the desmosomal protein DP. DP is required for the localization of Ect2 to keratinocyte desmosomes and cardiac intercalated discs in vitro and in vivo, where it maintains active RhoA (Rho-GTP) at the membrane. We demonstrate further that Ect2 activity is supported by PKC in a DP-dependent manner in cardiac myocytes. Finally, a truncated form of DP expressed in patients with Carvajal syndrome associated with severe cardiocutaneous defects is impaired in its ability to bind and localize Ect2 to cell junctions in cardiomyocytes and keratinocytes isolated from patients. Our findings delineate an important relationship between a component of the desmosome and a critical regulator of actin cytoskeletal remodeling that could have widespread implications for understanding cardiac and cutaneous health and disease pathogenesis.

## Introduction

Cell-cell junctions are vital sensors and integrators of mechanical forces experienced on the molecular, cellular and tissue scale. In addition to extrinsic forces experienced by the tissue, cell-cell junctions are also responsible for mediating intrinsic forces generated by cells of the tissue themselves, which are generally tensile forces that are generated when adhesion is coupled to the contractile actomyosin network [1-3]. One of the best characterized modulators of actomyosin contractility is the RhoA GTPase signaling pathway [4, 5]. Activation of RhoA leads to the direct modulation of its various downstream effectors, which have the cumulative outcome of promoting actin polymerization and myosin contractility. The importance of RhoA as a crucial regulator of junctional integrity and cellular contractility is now well-established [6-8]. Properly tuned RhoA activity is critical to maintain a balance of contractile activity at the cell cortex necessary for establishment and maintenance of polarized epithelial cell functions and interfering with Rho signaling is associated with disease pathogenesis [9].

Given the importance of RhoA in epithelia, it is surprising how little is known about junctional RhoA or the upstream factors that control its spatiotemporal activation in cardiac cell-cell junctions. Further, the extent to which these pathways are shared with those in epithelia are unknown. Unlike the multinucleated syncytium of skeletal muscle, cardiomyocytes are organized as discrete rod-shaped units that interconnect along their longitudinal axis via highly specialized intercellular junctions, referred to as intercalated discs (IDs) [10]. IDs represent the primary site of intercellular adhesion and electrochemical coupling between cardiomyocytes and can carry out these functions due to their unique junctional composition. Within IDs, components of adherens junctions (AJs), desmosomes and gap junctions are found in close proximity to each other and are able to form hybrid junctions that have been termed “area composita”, in which components of desmosomes and AJs are intermixed [11]. This is particularly important as IDs represent the primary site of force transmission between cardiomyocytes whereby the contractile forces generated by sarcomeres are transmitted to neighboring cardiomyocytes via these specialized junctions [10].

RhoA signaling has emerged as a somewhat controversial player in cardiac pathology [12]; however, despite the importance attributed to this contractile signaling modulator in epithelia its role in the ID is understudied. This is particularly notable given that intercellular junctions are major targets in disorders involving both skin and heart, including arrhythmogenic cardiomyopathy and related syndromes like Carvajal disease. 50% of patients presenting with these disorders harbor pathogenic variants of molecules in cadherin based intercellular junctions called desmosomes, which physically and functionally network with AJs. While heart and skin share the property of being mechanically stressed tissues, the underlying mechanisms linking specific variants to the heterogeneous phenotypes in these disorders is unknown.

Our lab previously reported that desmosome components play an important role along with their attached intermediate filaments (IF) in balancing the contractile activity mediated by AJs and their associated cortical actin [13, 14]. One such component is the cytolinker protein Desmoplakin (DP), which functions to tether IF to sites of cell-cell adhesion, providing mechanical integrity to the tissues in which it resides. Expression of a mutant version of DP lacking its IF binding domains (DPNTP) resulted in a depletion of actin dependent cellular forces that were associated with a concurrent reduction of active RhoA at epithelial cell junctions [13]. This finding, along with previous evidence that Plakophilin 2, a desmosomal component that is vital for DP’s localization to cell junctions, is involved in regulating RhoA GTPase junctional signaling [14], led us to hypothesize that DP could be acting as a scaffold for regulators of junctional RhoA GTPase signaling.

The class of proteins responsible for RhoA activation are the guanine nucleotide exchange factors (GEFs) that catalyze the exchange of GDP for GTP on Rho proteins, resulting in the activation of its GTPase function [15]. As such, the subcellular localization of specific GEFs is an important determinant of Rho GTPase activation at corresponding sites within a cell. The RhoGEF Ect2 has previously been documented to be localized to cell-cell junctions of interphase cells at steady-state in models of simple epithelia [16]. In simple epithelia, Ect2’s junctional localization was found to be dependent upon the AJ protein α-Catenin and once present at cell borders, forms a complex with α-Catenin and its binding partner E-Cadherin [16]. The junctional presence of Ect2 was further demonstrated to be crucial for the maintenance of an active pool of RhoA at sites of cell-cell adhesion and the promotion of junctional tension. However, it has yet to be established if Ect2 can be found at cellular junctions in more complex tissues such as the stratified epithelium of the epidermis or the myocardium and to what extent it could be playing a role in modulating the junctional tension and cellular forces within these tissues.

In this study, we identify the RhoGEF Ect2 as a previously unappreciated component of the cardiac intercalated disc (ID) and binding partner of the desmosomal protein DP. We further demonstrate that DP is required for the localization of Ect2 to cardiac junctions in both in vitro and in vivo models, independent of the AJ machinery that it utilizes in simple epithelial models. Depletion of either DP or Ect2 was found to be sufficient to significantly perturb the maintenance of an active pool of RhoA that is present at steady-state at cardiac cell junctions. Finally, we demonstrate that a truncated form of DP harboring a patient mutation that results in severe cardiocutaneous defects is impaired in its ability to bind and localize Ect2 to cell junctions in cardiomyocytes and keratinocytes isolated from patients. Our findings delineate an important regulatory relationship between a component of the desmosome and a critical regulator of cytoskeletal remodeling that could have widespread implications towards our understanding of cardiac and cutaneous health and disease progression.

## Results

### Ect2 is a novel binding partner of Desmoplakin and is a component of the cardiac intercalated disc

We previously showed that expression of DP mutants that uncouple the IF network from the desmosomal complex impairs the junctional distribution and activation of the small GTPase RhoA [13]. As it is well-established that spatiotemporal control of RhoA activation is heavily dependent upon the localization of GEFs to specific sites within the cell, we hypothesized that DP acts as a scaffold for a RhoGEF at cardiac cell junctions. Previous studies identified the RhoGEF Ect2 as a critical regulator of junctional RhoA GTPase activation and junctional tension in simple epithelial cells and as such, Ect2 is a plausible candidate regulator of RhoA activity in cardiac junctions.

We first tested for the presence of Ect2 at cardiac cell junctions utilizing the well-established primary cell culture model of neonatal rat ventricular cardiomyocytes (NRVCMs). Immunostaining of Ect2 and DP carried out on freshly isolated NRVCMs, followed by structured illumination microscopy (SIM) imaging showed that Ect2 is indeed found to be enriched at cardiac cell junctions (Figure 1A; Supplementary Figure 1A). Additionally, when overlaid with DP immunostaining at corresponding junctions, Ect2 displayed strong co-localization with DP (Figure 1A; Supplementary Figure 1A). Having confirmed the presence of Ect2 at cardiac junctions, we next addressed whether this localization is conserved in vivo. To this end, we carried out immunostaining of Ect2 and DP on murine and human cardiac tissue and observed a robust Ect2 positive signal co-localizing with DP at murine and human IDs (Figure 1B, 1B’; Supplementary Figure 1 B,C). Immunoprecipitation of DP followed by blot back with an Ect2 antibody in human keratinocytes showed that Ect2 was specifically pulled down with DP, compared with the IgG control (Supplementary Figure 1D). To corroborate Ect2 as a possible interacting partner of DP at the plasma membrane, we carried out proximity ligation assay (PLA) analysis of DP and Ect2 in NRVCMs. This method allows for the detection of proteins that are in close proximity with one another (40-100 nm) and is frequently utilized to detect in situ protein-protein interactions. NRVCMs were treated with adenovirus harboring either control (non-targeting) or DP KD shRNA constructs followed by PLA analysis. Control NRVCMs showed a robust DP-Ect2 PLA signal that significantly decreased in DP KD cells, indicating that this signal was specific to the DP-Ect2 antibody pairing (Figure 1C & 1C’).

**Figure 1:**
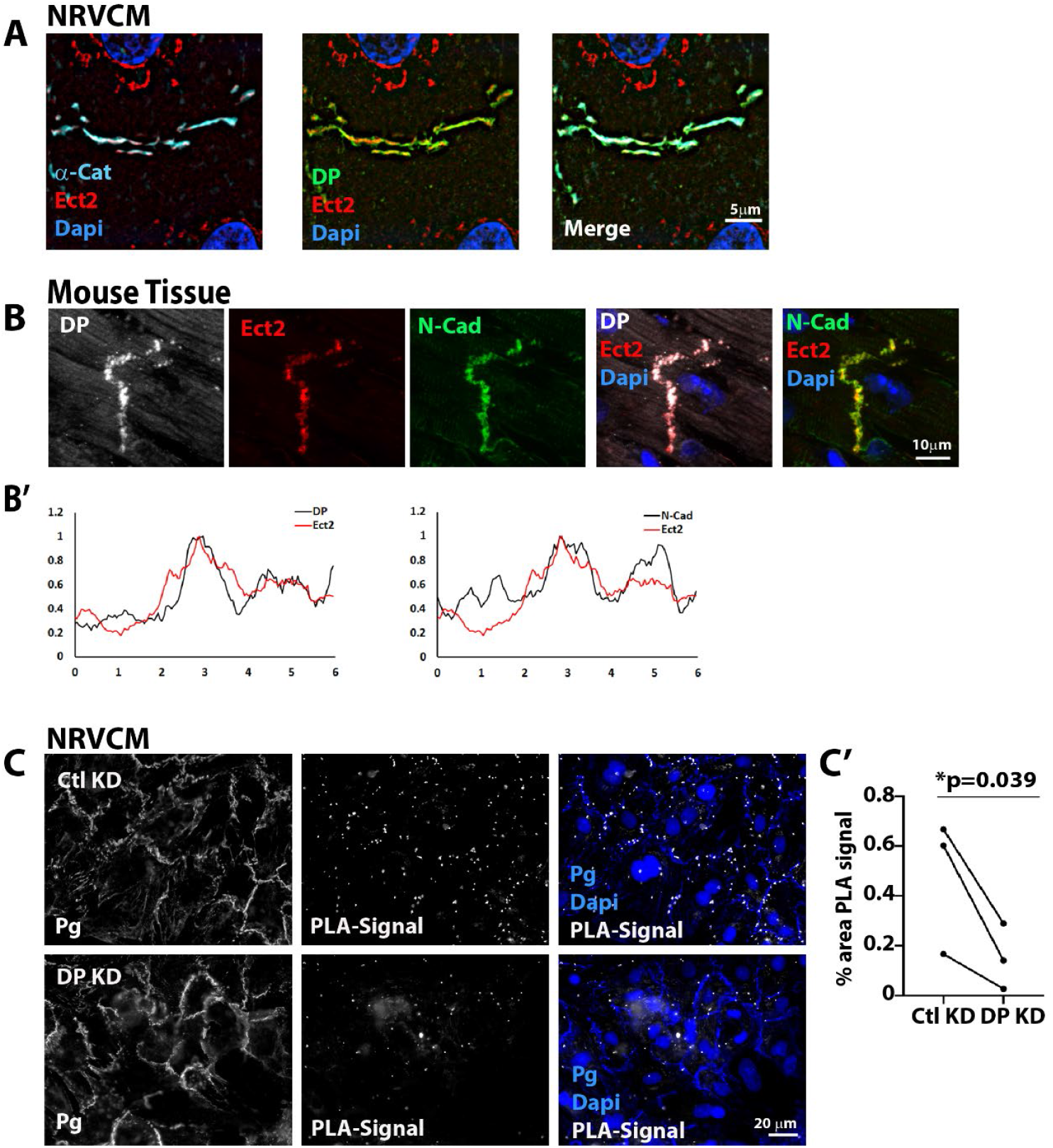
Desmoplakin and the RhoGEF Ect2 co-localize at the intercalated disc. (A) Isolated cardiac myocytes (NRVCM) were prepared for immunofluorescence and labeled for Desmoplakin (DP), Ect2 and α-Catenin and imaged using structured illumination microscopy (SIM). Orthogonal views can be found in Supplemental Figure 1A. Scale bar = 5 μm. (B) Sections from control mouse hearts were fixed and stained for DP, Ect2 and N-Cadherin (N-Cad) and imaged using the AxioVison Z1 system (Carl Zeiss) with Apotome slide module. Scale bar = 10μm (B’) Fluorescence intensity for DP, Ect2 and N-Cad was quantified in a line scan across the ID, showing greater correspondence between DP and Ect2 than N-Cadherin and Ect2. (C) Proximity ligation assay (PLA) was performed on NRVCM in Control (Ctl) and DP knockdown (KD) conditions using antibodies directed against DP and Ect2. Scale bar = 20μm. (C’) Quantification of each of three paired experiments was performed, showing significantly more PLA spots in Ctl KD conditions, indicating close proximity of DP and Ect2 (p=0.039). Dapi is used to stain nuclei in A, B, C.

These data indicate that Ect2 is a previously unappreciated binding partner of DP in cardiac and epithelial cells and localizes with DP at cell-cell junctions. Furthermore, they also show that Ect2 is a novel component of the ID complex. Given the importance of spatial distribution of GEFs in localizing and activating RhoA, we carried out further experiments to elucidate the role played by the DP-Ect2 complex in cardiac function and homeostasis.

### DP is required for the localization of Ect2 to cardiac cell junctions

Towards understanding the importance of DP for Ect2 localization, we first asked whether DP is required for Ect2’s recruitment to cardiac cell junctions. NRVCMs were treated with adenovirus harboring control, DP KD, or Ect2 KD shRNA constructs followed by immunofluorescence staining of DP and Ect2. The desmosomal component Plakoglobin (Pg) was also labeled as a marker of cell junctions as we have previously shown that Pg junctional localization is not significantly affected by depleting DP in cardiac cells [17]. Efficacy of the shRNA directed toward DP and Ect2 was confirmed by the loss of immunofluorescence signal (Figure 2A).

**Figure 2:**
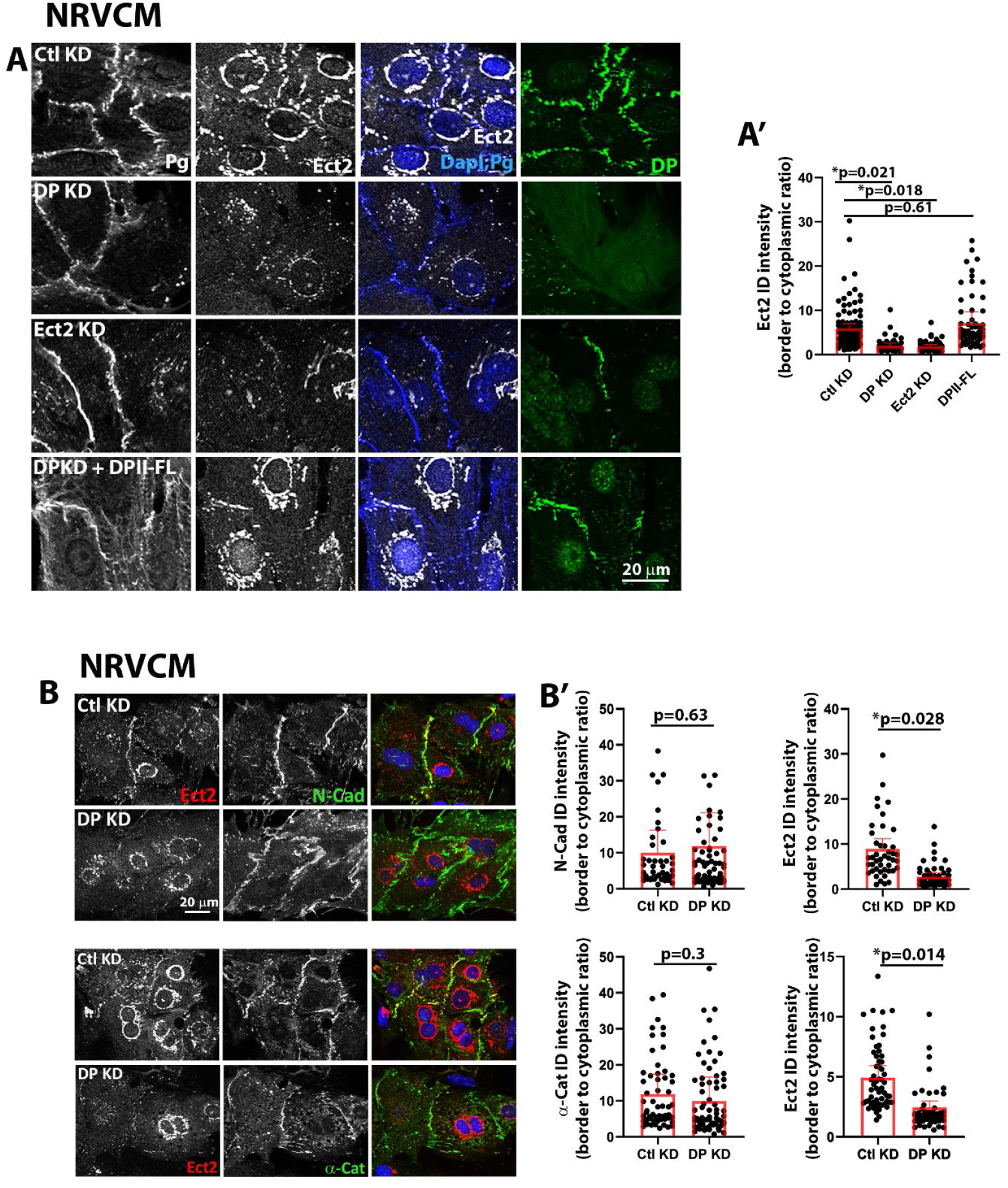
DP is required for the localization of Ect2 to cardiac cell junctions in vitro. (A) Isolated cardiac myocytes (NRVCM) were treated with adenovirus encoding scrambled control (Ctl), DP or Ect2 knockdown (KD) oligonucleotides, and then stained for DP and Ect2 as well as plakoglobin (Pg) to mark cell-cell borders. In addition, DP KD was rescued by expression of full-length Desmoplakin II (DPII-FL). (A’) Quantification of fluorescence intensity at the cell-cell junctions showed that Ect2 junctional staining was lost in the absence of DP and could be rescued by ectopic DPII expression. (B) NRVCM were treated with adenovirus encoding scrambled control (Ctl) or DP KD oligonucleotides and stained for Ect2 and the adherens junction proteins N-Cadherin (N-Cad) or α-Catenin (α-Cat). (B’) Quantification of fluorescence intensity at the cell-cell junctions showed that neither N-Cadherin or α-Catenin were significantly changed with DP KD, but Ect2 border intensity significantly decreased (p<0.05). All statistical tests were performed on the means of three independent experiments.

Depletion of DP in NRVCMs resulted in an almost complete abolishment of Ect2 signal at cardiac cell junctions (Figure 2A). This DP dependent decrease of junctional Ect2 signal was found to be significant by quantifying Ect2 border signal (border to cytoplasmic ratio, border position indicated by Pg) (Figure 2A’). We also noted a reduction in DP at borders in response to Ect2 KD, suggesting a mutual dependence on protein localization and/or stability. There is also an Ect2 fluorescent signal marking the periphery of the nucleus, which was only partially affected by Ect2 KD shRNA, suggesting the possibility that this signal is at least in part non-specific. To verify that loss of junctional Ect2 is specifically due to DP KD, we re-expressed exogenous DP by introducing an adenovirus harboring a DPII-GFP construct in the background of DP KD. DPII is a naturally occurring isoform of DP that lacks a portion of its central rod domain (6.8 vs 8.6 kb), which allows us to overcome the technical limits in size for adenoviral packaging and transduction. DPII expression in a DP deficient background was sufficient to significantly restore Ect2’s localization to cardiac cell junctions (Figure 2A & 2A’).

A previous study in simple epithelial cells reported that Ect2 forms a complex with the AJ components E-Cadherin and α-Catenin and co-localizes with AJs [16]. Furthermore, the disruption of AJs via KD of α-Catenin was sufficient to disrupt Ect2’s localization to epithelial borders. To determine whether the failure of Ect2 to localize to cell borders under DP KD conditions was due to concurrent perturbations to AJs, we sought to assess the status of the cardiac AJ components, α-Catenin and N-Cadherin via immunofluorescence analysis in a DP deficient background. Ect2, N-Cadherin and α-Catenin were present at cell borders in control NRVCMs as expected, however even though DP KD led to a significant reduction in junctional Ect2, there were no significant changes in N-Cadherin or α-Catenin at corresponding junctions where Ect2 was present (Figure 2B & 2B’).

We then asked whether DP dependent junctional localization of Ect2 is conserved in vivo, using a previously reported murine cardiac conditional DP knockout (*Dsp* cKO) model [18]. This model was generated by crossing *Ds*p-floxed mice (*Dsp*^flox/flox^) with the well-established cardiomyocyte-specific ventricular myosin light chain-2-Cre recombinase (MLC2v-Cre) knock-in mice. Cardiac sections from 8 week old wildtype (WT) and *Dsp* cKO mice were immunostained with antibodies directed against DP, Ect2 and Pg, as a marker of IDs. As previously reported, DP staining was almost completely abolished in *Dsp* cKO hearts with occasional retention of DP in a small number of IDs (Figure 3A). *Dsp* cKO cardiac sections also displayed a significant reduction in Ect2 signal at IDs (Figure 3A & 3A’). Consistent with our in vitro experiments, co-staining of Ect2 with either N-Cadherin or α-Catenin in *Dsp* cKO hearts showed that junctional Ect2 signal was depleted even though ID intensity of N-Cadherin or α-Catenin were not significantly affected (Figure 3B, 3B’, 3C, 3C’). These data indicate that DP is a specific regulator of Ect2 junctional localization in cardiomyocytes in culture and adult mouse cardiac tissue.

**Figure 3:**
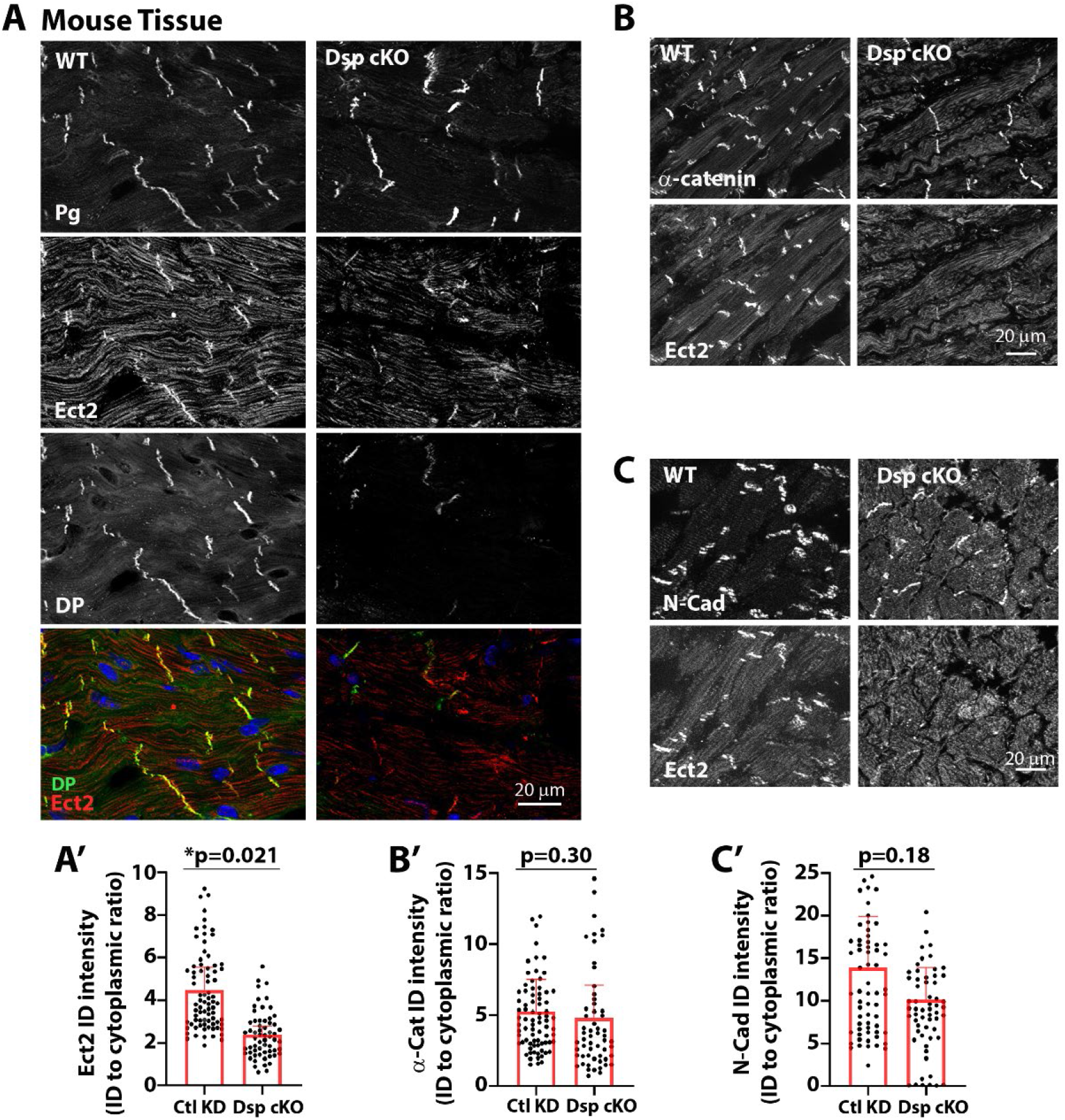
DP is required for the localization of Ect2 to cardiac cell junctions in vivo. (A) Sections from control and *Dsp* cKO mouse hearts were fixed and triple-labeled for DP, Ect2 as well as Pg to mark intercalated disks (IDs). (A’) Quantification of fluorescence intensity at IDs showed a significant decrease in Ect2 at *Dsp* cKO IDs (p=0.021). (B) Sections from control and *Dsp* cKO mouse hearts were fixed and double-labeled for α-Catenin and Ect2. (B’) Quantification of fluorescence intensity at IDs showed no significant decrease in α-Catenin at *Dsp* cKO IDs although Ect2 was decreased. (C) Sections from control and *Dsp* cKO mouse hearts were fixed and double-labeled for N-Cadherin and Ect2. (C’) Quantification of fluorescence intensity at IDs showed no significant decrease in N-Cadherin at *Dsp* cKO IDs although Ect2 was decreased. Scale bars = 20μm. All statistical tests were performed on the means of three independent experiments.

### DP loss results in reduction of activated RhoA at cardiac cell junctions

A role for RhoA signaling in the regulation of cardiac development and homeostasis is well documented, if somewhat controversial [12, 19, 20]. However, the role and regulation of junctional specific RhoA in cardiac systems has gone largely unexplored. To assess the status of RhoA and its activation in cardiac junctions, we employed the well-established TCA fixation method that allows for the visualization of junctional specific endogenous RhoA by immunofluorescence [21, 22]. Utilizing this fixation method, we assessed the distribution of active RhoA in NRVCMs with an antibody that specifically recognizes the GTP bound active form of RhoA (Rho-GTP). In addition to the expected cytoplasmic distribution of RhoA, we detected a concentration of RhoA-GTP signal along the cell-cell junctions of control NRVCMs as visualized by co-staining with Pg (Figure 4A). Furthermore, DP or Ect2 KD resulted in a significant reduction of junctionally localized RhoA-GTP, confirming the role of these proteins in the maintenance of the junctional specific pool of active RhoA in cardiac cells (Figure 4A, 4A’). Re-expression of full length DPII in a DP deficient background was sufficient to restore the RhoA-GTP signal back to that of control levels (Figure 4A, 4A’).

**Figure 4:**
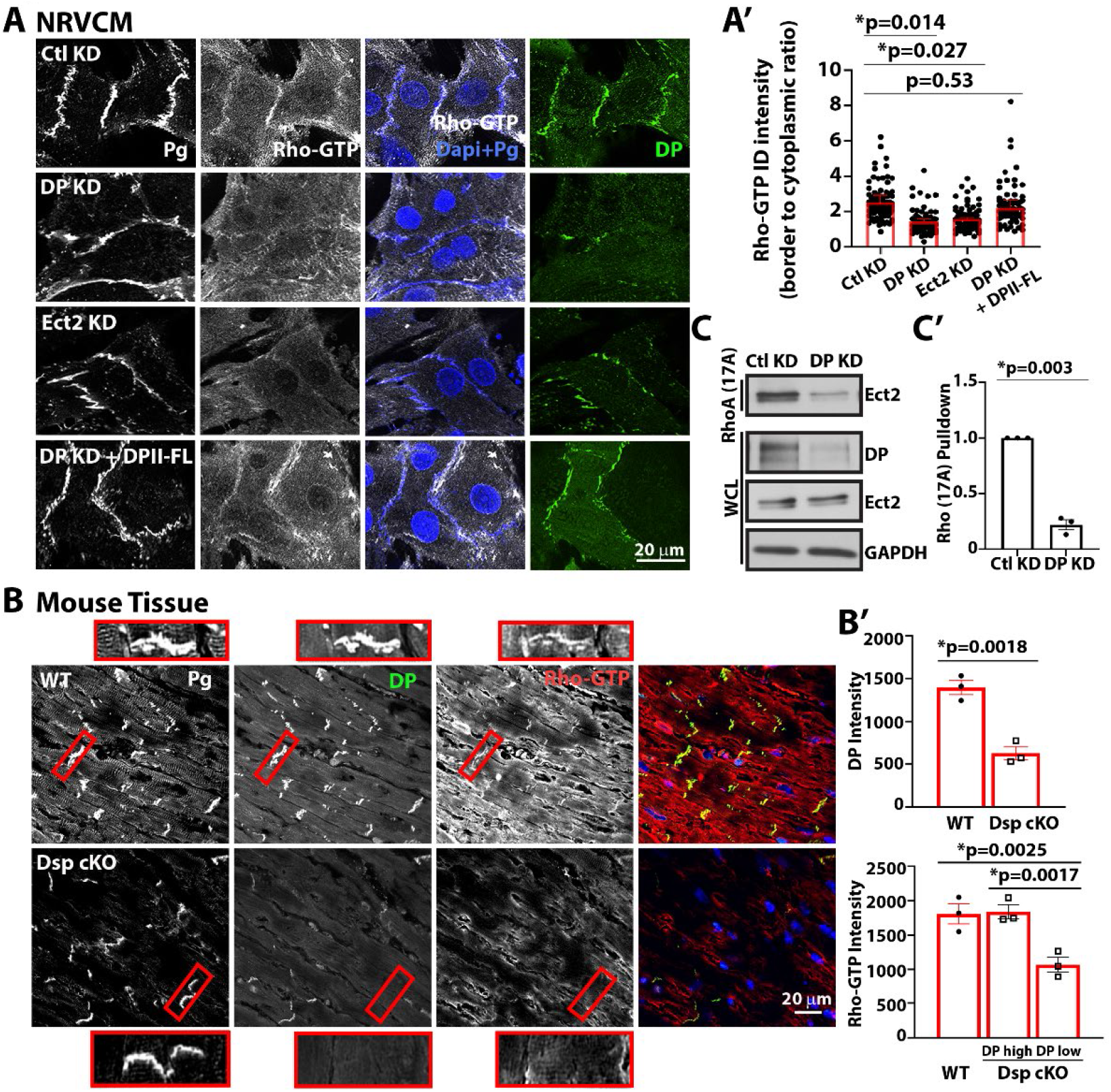
Active RhoA at cardiac cell junctions requires the expression of DP and colocalizes with DP at IDs. A) Isolated cardiac myocytes (NRVCM) were treated with scrambled control (Ctl), DP or Ect2 knockdown (KD) oligonucleotides, and then stained for plakoglobin (Pg) to mark cell-cell borders, as well an antibody that recognizes active RhoA (Rho-GTP). Active RhoA cell junction localization is lost upon either DP or Ect2 silencing and its border localization is restored by ectopic expression of DPII-FL, quantified in (A’). (B) Sections from control and *Dsp* cKO mouse hearts were fixed and triple-labeled for Pg to mark IDs, DP and active RhoA. (B’) Quantification of DP border intensity in IDs of the WT vs *Dsp* cKO animals and Rho-GTP intensity at IDs of the WT and *Dsp* cKO animals showed a significant decrease in active RhoA at IDs with low DP localization compared to WT (p=0.0025) or IDs that retain DP (DP high IDs, p=0.0017). (C) Lysates from Ctl or DP KD HL-1 cardiac myocytes were subjected to RhoA (17A) pull down to isolate active GEFs and lysates were probed for Ect2. Whole cell lysates (WCL) were probed for DP and Ect2 as well as GAPDH as a loading control. (C’) Quantification of blots demonstrated a significant decrease in active Ect2 in DP KD lysates (p=0.003). Scale bars = 20μm. All statistical tests were performed on the means of three independent experiments.

We also tested if these observations held true in vivo, utilizing the murine model of cardiac *Dsp* cKO. Immunofluorescence analysis confirmed that RhoA-GTP signal is concentrated at mature IDs of WT mouse heart sections (Figure 4B & insets). Supporting our in vitro findings, *Dsp* cKO cardiac sections displayed a significant reduction of ID specific RhoA-GTP signal at IDs where DP signal was also absent (Figure 4B & insets, 4B’; Supplementary Figure 2). These data demonstrate that DP is required for the maintenance of an active pool of RhoA-GTP at cardiac cell junctions. Having shown that independent KD of either DP or Ect2 was sufficient to cause a reduction of activated RhoA at cardiac cell junctions, we next assessed if DP controls Ect2’s catalytic GEF function towards RhoA. To assess this, we employed an assay designed to precipitate the active pool of GEFs from cell lysates. This technique takes advantage of a single amino acid mutation in RhoA (G17A) that is stabilized in a nucleotide-free state, which has a high affinity towards active RhoA-specific GEFs [23]. Utilizing this assay in HL-1 cardiac cells, we observed that DP KD resulted in a significant reduction of Ect2 precipitated by RhoA (17A) beads compared to control lysates (Figure 4C, 4C’). This result confirms that DP is an important regulator of Ect2’s ability to catalyze the exchange of GDP for GTP on RhoA.

### DP regulates Ect2 activity via PKC mediated phosphorylation

We next set out to determine the mechanism by which DP regulates activation of Ect2. It was previously reported that protein kinase C (PKC)-dependent phosphorylation of Ect2 upregulates its activity [24, 25]. Further, our lab previously showed that DP forms a complex with PKCα and its desmosomal binding partner, Plakophilin-2 [26]. As such, we tested the hypothesis that DP could be acting as a scaffold for PKC at cardiac cell junctions to allow for phosphorylation and activation of Ect2.

We first assessed if Ect2’s activation state in cardiac cells is responsive to alterations in PKC kinase activity levels. To do so, we leveraged the aforementioned Rho (G17A) pulldown assay to determine Ect2’s activation levels in the presence of pharmacological modulation of PKC activity. Treatment of HL-1 cells with the broad-spectrum PKC inhibitor, BIM1, led to a significant decrease in Ect2 activity (Figure 5A, 5A’). Conversely, following treatment with the well-established PKC activator, PMA, Ect2 activation status was significantly upregulated in HL-1 cells, suggesting that Ect2 activation can be modulated by PKC kinase activity.

**Figure 5:**
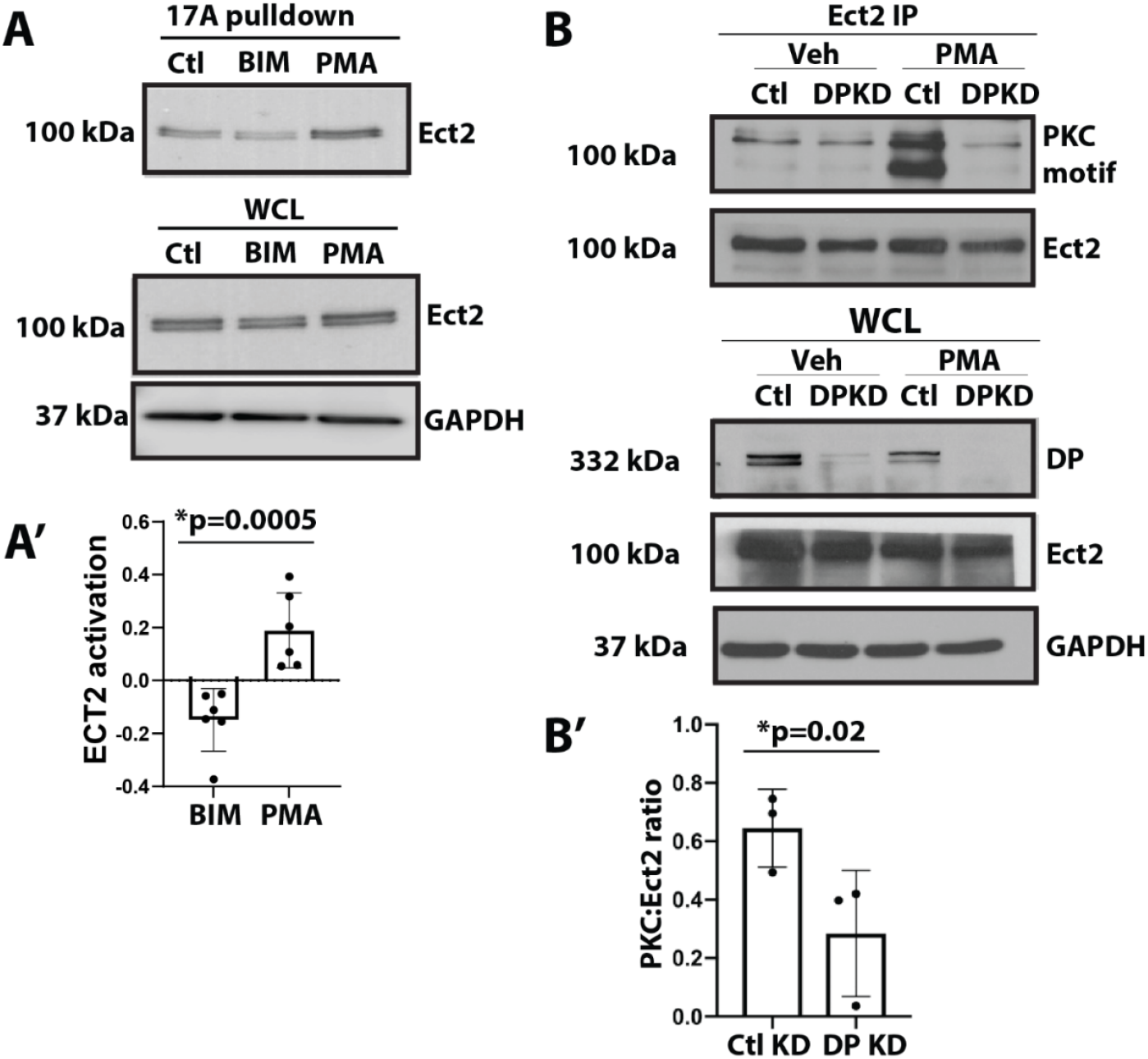
DP regulates Ect2 activity via PKC mediated phosphorylation. (A) Lysates from Ctl, BIM or PMA treated HL-1 cells were subjected to RhoA (17A) pull down to isolate active GEFs and probed for Ect2. Whole cell lysates (WCL) were probed for Ect2 as well as GAPDH as a loading control. (A’) Quantification of blots demonstrated significantly more active Ect2 in cells in which PKC has been activated (PMA), compared to BIM-treated cells in which PKC is inhibited (p=0.0005). Data are from 5 independent experiments. (B) Ect2 was immunoprecipitated from Ctl or DP KD HL-1 cells treated with vehicle or PMA-treated and blotted back with an antibody recognizing PKC-dependent phosphorylation or an antibody against Ect2. WCLs were probed for DP and Ect2 as well as GAPDH as a loading control. (B’) Significantly less PKC-dependent phosphorylation was detected on Ect2 in DP KD cells (p=0.02). Data are from three independent experiments.

To determine if DP plays a role in regulating PKC-mediated phosphorylation of Ect2, we next carried out immunoprecipitation (IP) of endogenous Ect2 in control and DP KD NRVCMs treated with either vehicle (DMSO) or PMA. This was followed by blot back for Ect2, as well as with an antibody that recognizes phosphorylated PKC substrates. Immunoblotting for PKC phosphorylated substrates in vehicle treated control KD NRVCMs showed positive signal at the molecular weight corresponding to Ect2 (Figure 5B). Importantly, while control KD samples treated with PMA displayed a robust elevation of the PKC substrate motif signal, there was no apparent change in PMA treated DP KD samples compared to control (Figure 5B, 5B’). These findings indicate that DP is required for PKC-mediated phosphorylation of Ect2, possibly acting as a scaffold for these proteins.

### Ect2 junctional localization is disrupted in cardiac and epidermal cells from cardiocutaneous disease patients

Having delineated the role that DP plays in regulating the junctional localization and activity of Ect2 in cardiac models, we next asked whether the functional relationship between DP and Ect2 could have consequences for disorders associated with mutations in the *DSP* gene. Inherited mutations in *DSP* have been found to be a frequent causative factor underlying myocardial disease including Arrhythmogenic Cardiomyopathy (AC), dilated cardiomyopathy and myocarditis [27-29]. Variants include Carvajal syndrome, a familial cardiocutaneous disorder defined by the phenotypic presentation of woolly hair, palmoplantar keratoderma and cardiac disease resulting from mutations in the *DSP* gene [30]. Given the reported loss of DP localization to IDs of Carvajal patients, and our demonstration that DP was required for Ect2 localization at murine IDs (Figure 3), we asked whether Ect2 is also perturbed in tissue from a Carvajal patient [30]. Utilizing N-Cadherin signal as a marker of IDs, we observed that ID localization of Ect2 is completely lost in cardiac sections from the Carvajal Syndrome patient when compared to healthy control tissue (Figure 6A).

**Figure 6:**
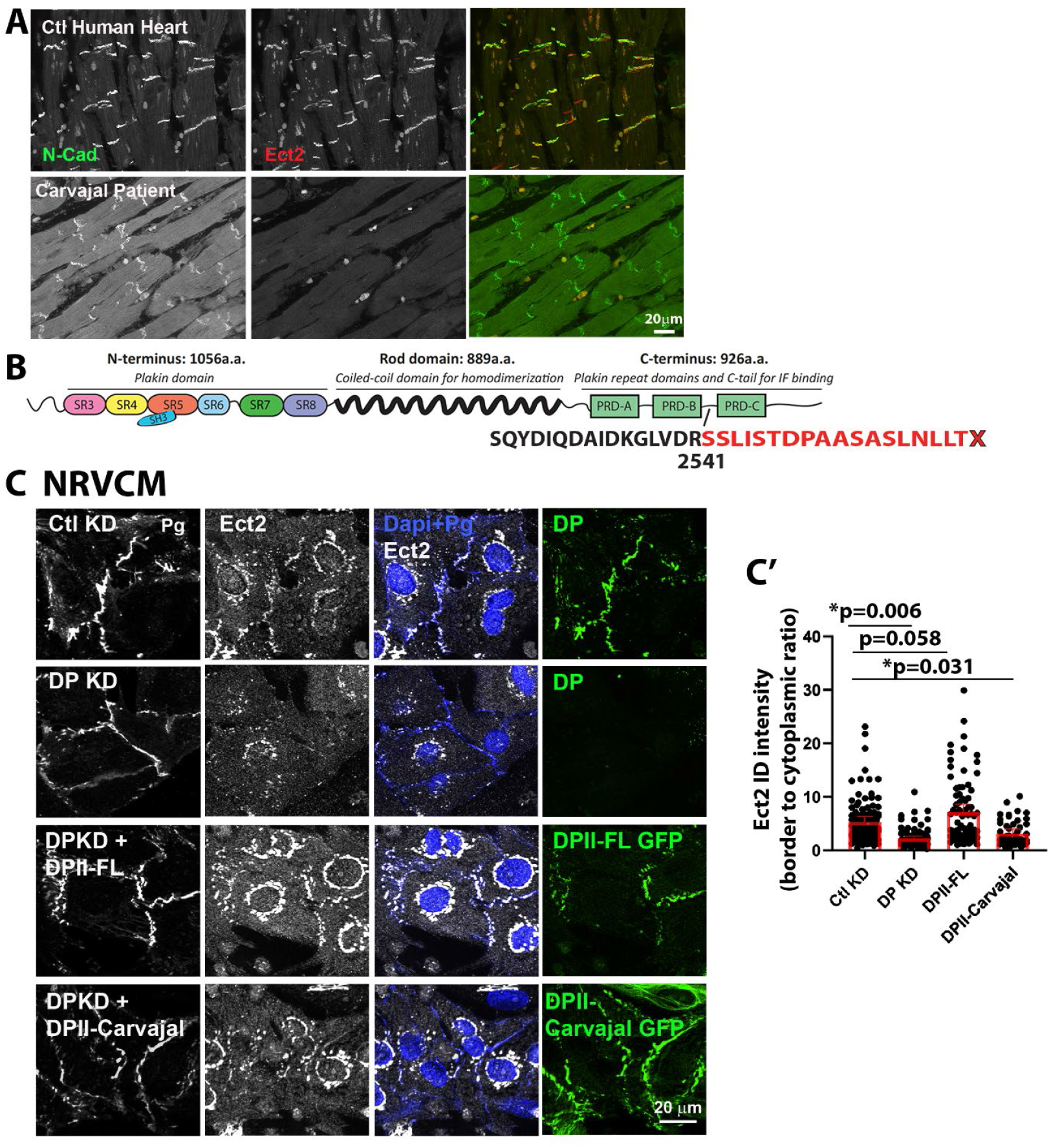
Ect2 junctional localization is disrupted in cardiac tissues from Carvajal patients. (A) Sections from control (Ctl) or human Carvajal patient heart tissue were fixed and stained for Ect2 as well as N-Cadherin to mark IDs. Ect2 staining at IDs was present in Ctl but not Carvajal samples. Scale bar = 20μm. (B) Structure of DP from Carvajal patients, compared with WT DP. Note that the map shows the location of the point mutation in DPI (aa2541) while the engineered constructs are in DPII (aa 1941). (C) Isolated NRVCM were treated with adenovirus encoding scrambled control (Ctl) or DP KD oligonucleotides and rescued with either wild type DPII-FL or DPII-Carvajal constructs and stained for Ect2 and DP. (C’) Quantification of fluorescence intensity of Ect2 at IDs showed that while Ect2 was observed at junctions in DPII-FL expressing cells, the Carvajal mutant was unable to restore Ect2 at IDs. Statistical tests were performed on the means of three independent experiments.

A Carvajal disease patient reported in the literature harbored a single point mutation in the *DSP* gene that resulted in a premature stop codon and truncation of the third plakin repeat domain within the C-terminus of DP (Figure 6B) [31]. This patient presented with dilated cardiomyopathy, a form of cardiac disease characterized by an enlargement of the ventricular chamber and thinning of ventricular walls, along with cutaneous symptoms [30]. Given the presentation of both cardiac and cutaneous phenotypes in this patient, we asked whether the junctional localization of Ect2 is impaired in keratinocytes isolated from this patient. To test this, control immortalized human keratinocytes (IHEKs) and immortalized keratinocytes isolated from the Carvajal syndrome patient (JD1) were cultured for 2 days in high calcium media to promote differentiation and desmosome maturation. In contrast to the cardiac sections from this patient, immunofluorescence analysis of these samples showed that DP was still able to localize to cell-cell junctions, suggesting a possible difference in the incorporation of this mutant protein into the cell junctions of these tissue types (Supplementary Figure 3A). We also observed that Ect2 co-localized at keratinocyte cell junctions in IHEKs; however, the Ect2 junctional signal was significantly reduced in JD1 cells (Supplementary Figure 3B).

To interrogate further the functional role that this DP mutant could be playing in the regulation of Ect2, we engineered the same point mutation that results in the truncated form of DP in the Carvajal syndrome patient into the DPII-GFP construct and inserted it into our adenoviral vector (Figure 6B). Interestingly, immunostaining of the Carvajal mutant DPII (DPII-Carvajal) in NRVCMs showed that this mutant form of DP was still able to localize to cardiac cell junctions, similar to the observation in JD1 cells (Figure 6C). This could suggest that loss of DP from the Carvajal patient heart could be due to the chronic effect of compromised cardiac function over time. Interestingly, immunostaining of Ect2 revealed a significant decrease of junctional Ect2 in DPII-Carvajal cells compared to WT DPII samples (Figure 6C, 6C’). To corroborate that the Carvajal mutation compromises the ability of DP to recruit Ect2, a GFP peptide trap was used to pull down the GFP tagged WT DPII and DPII-Carvajal fusion proteins. Ect2 was pulled down along with WT DPII-GFP in the GFP trap, but was not detected in the GFP only control or in the Carvajal-GFP pull downs (Supplementary Figure 3C). These data suggest that a portion of the DP C-terminus is required for the localization of Ect2 to cardiac cell junctions and could represent a binding site on DP for Ect2, necessary for RhoA localization and junctional activity.

## DISCUSSION

In this study, we identified the RhoGEF Ect2 as a novel component of the cardiac ID and keratinocyte desmosomes, which is required for the maintenance of an active pool of RhoA at IDs. Previous work has identified several RhoGEFs important for cardiac development and homeostasis [32-34], including sarcomeric-localized RhoGEFs such as p63RhoGEF [35] and Obscurin [33]. However, despite the established importance of subcellular positioning of RhoA activators, to our knowledge, this is the first report of a RhoA GEF that specifically localized to ID structures.

Rho signaling through other molecular complexes present at the ID have been reported to activate downstream pathways through kinases and transcription factors including ROCK, YAP, MRTF-A and PLCε, to regulate cardiac development and pathophysiology [20]. For instance, the protein Myozap was identified as a component of the ID that directly binds to DP and the myosin phosphatase-RhoA interacting protein (MRIP), a negative regulator of Rho activity. Myozap activates MRTF/SRF-dependent signaling in a Rho-dependent fashion, and its loss has been linked to contractile dysfunction and cardiomyopathy [36]. More recently, the GTPase Rnd1 was identified as a Myozap partner that promotes SRF signaling in a stretch-sensitive manner to regulate cell proliferation and hypertrophy [37]. The AJ and ID protein α-Catenin, which serves a cytoskeletal adapter role similar to DP in the AJ, regulates ID maturation and actomyosin contractility in a mechanosensitive fashion, which, in turn, controls the cytoplasmic to nuclear distribution of the transcription factor Yap to regulate cell proliferation [38]

The identification of Ect2 as another DP binding partner supports the existence of a more direct pathway to RhoA activation that could functionally complement mechanisms driven by Myozap. For instance, while Myozap activates long term responses through SRF transcription, it is possible that more direct coupling of desmosomes to RhoA activation through Ect2 could lead to rapid changes in mechanical signaling. The desmosome-IF network plays a crucial role in balancing the forces of mechanical stress in many systems, allowing excess contractile activity to be offloaded [2]. This capacity is particularly important in the ID, which facilitates force transmission and transduction between cells and across the myocardium [39]. Indeed, using a desmoglein-2 (DSG-2) force sensor, high tensile loading was measured during contraction of human cardiac muscle cells, raising the possibility that desmosomes can directly sense and respond to mechanical stress [40]. The DP-Ect2 module may serve as a sensor as a part of a relay that helps synchronize these signals across the tissue in rapid fashion.

Here we show that DP not only binds and co-localizes with Ect2, but also regulates Ect2 activity through PKC, a family of kinases that play critical roles in cardiac remodeling including during cardiac hypertrophy [41]. We previously demonstrated that DP exists in a complex with PKCα through its interactions with the armadillo protein Plakophilin 2, on which it depends for its efficient localization to intercellular junctions [26]. Together, these observations suggest that desmosomes govern the localization and activity of a RhoA signaling module in the ID. The known dependence of PKC activity on the mechanical environment [42, 43], provides another mechanism to couple this module to changes in tissue mechanics during development and disease pathogenesis.

Several lines of evidence have linked aberrant Rho signaling to cardiac disease. In an animal model, a dominant active version of the downstream mediator ROCK was demonstrated to recapitulate fibrosis associated with heart failure in humans, where active ROCK1 accumulates [44]. Consistent with these results, ROCK1-/-mice exhibited reduced fibrosis in the myocardium associated with reduced expression of extracellular matrix proteins and fibrogenic cytokines [45]. In addition to a role for dysregulated RhoA signaling in cardiac disease, Rho/ROCK signaling contributes to cell fate decisions including cardiomyocyte identity through transcription of MRTF/SRF targets. Interfering with this pathway can prime cells to switch to an adipocyte lineage during cardiomyocyte differentiation [46].

We showed that Ect2 binding on DP depends at least in part on a C-terminal region previously shown to comprise a portion of the IF binding domain, followed by 68 residues that dictate the strength of IF binding through post-translational modifications [47-49]. This region is missing in a family with Carvajal disease, a syndrome associated with both cardiomyopathy and cutaneous symptoms including wooly hair [31]. We observed a partial loss of Ect2 associated with DP in keratinocytes isolated from the skin of a patient, and a loss of both DP and Ect2 at IDs in cardiac tissue from one of these patients. This raises the possibility that a progressive loss of junctional Ect2 function occurs as the concealed phase transitions to more advanced cardiomyopathy state. Additional functions of DP may also be lost during this transition, for instance proteome degradation control through loss of the cardiac constitutive photomorphogenesis 9 (COP9) signalosome subunits 3 or 6 at the desmosome [50] [51].

Finally, we previously demonstrated that epithelial sheets generated from Carvajal keratinocytes have reduced ability to withstand mechanical stress, consistent with impaired adhesive strength [52]. It will be interesting in the future to determine the extent to which DP-Ect-RhoA module works to convert mechanical cues into alterations in contractile signaling versus longer term transcriptional changes depending on actin remodeling, for instance through the SRF/MRTF pathway, which has been demonstrated to be important for cardiac homeostasis and epidermal differentiation [46, 53-56].

## Materials and Methods

### Isolation of NRVCMs, cell culture, mouse models and human tissues

Neonatal rat ventricular cardiomyocytes were isolated as described from one to three day old Sprague-Dawley rats (Charles River) [57]. Briefly, hearts were dissected from neonatal rats, enzymatically digested, and resuspended in M199 medium (Lonza) containing 10% FBS and supplemented with 15 μM vitamin B12. To deplete isolates of more rapidly adhering cardiac fibroblasts, cells were plated for 2h on plastic cell culture dishes. The remaining cell suspension, enriched for cardiac myocytes, was then filtered through a 40 μM filter, spun down, and resuspended in M199 medium containing 10% horse serum, 15 μM vitamin B12, and BrdU. Cells were then plated onto 6-well dishes or coverslips that had been coated with collagen type IV. Media was changed every 24-48 h with adenoviral infection being performed 48 h post isolation. These protocols were conducted with the approval of the Northwestern University Institutional Animal Care and Use Committee, and all animal care protocols conform to National Institutes of Health guidelines and the recommendations of the Panel on Euthanasia of the American Veterinary Medical Association.

The HL-1 cardiac myocyte cell line was maintained with Claycomb Medium (Sigma-Aldrich) supplemented with 10% FBS (Atlanta Biologicals), 0.1 mM norepinephrine, 2mM L-glutamine, and penicillin/streptomycin solution (Sigma-Aldrich). Cells were grown on plastic dishes precoated with a solution of FN-0.02% gelatin. The Carvajal keratinocyte line JD1 harbors DP with a deletion of G at position 7622 within the coding sequence of DP (position 7901 of Genbank/EMBL/DDBJ accession no. M77830). This mutation results in loss of the C-subdomain containing the plakin repeats and part of the upstream linker region and introduces 18 new amino acids downstream of the deletion. JD-1 keratinocytes were isolated and immortalized with an HPV16 plasmid (pJ45216) that has early region genes driven by MoMLV-LTR as previously described [58]. Normal adult keratinocytes immortalized with HPV16 E7 (IHEKs) were obtained from the Skin Biology and Diseases Resource Based Center at Northwestern University Feinberg School of Medicine. Cells were cultured in DME/Ham’s F12 (3:1) supplemented with 10% FCS, 4 mM glutamine, 0.4 μg/ml hydrocortisone, 0.1 nM cholera toxin, 5 μg/ml insulin, and 10 ng/ml EGF.

Primary normal human epidermal keratinocytes (NHEKs) were isolated from human foreskin as previously described and grown in M154 media supplemented with 0.07 mM CaCl_2_, human keratinocyte growth supplement (HKGS) and gentamicin/amphotericin B solution (Thermo Fisher Scientific) [59]. NHEKs were grown to confluency and switched to M154 media supplemented with HKGS, gentamicin/amphotericin B and 1.2 mM CaCl_2_ for 24 h-48 h.

Mouse tissues were obtained from animals in which the *Dsp* gene was deleted in the mouse myocardium by breeding *DP*-floxed mice with heterozygous ventricular myosin light chain-2 Cre (MLC2v^(cre+)^) mice. *Dsp* cKO mice and their control littermates were kept in a congenic C56Bl/6 background. Hearts obtained from 8-week old mice were snap frozen in liquid nitrogen and sectioned for immunofluorescence analysis.

Human tissues were obtained from Dr. Jeffrey Saffitz (Beth Israel Deaconess Medical Center, Boston, MA). De-identified control and Carvajal patient left ventricular myocardial tissue were obtained in paraffin embedded blocks. Tissue sections were processed for immunohistochemistry as described below.

### DNA constructs, siRNAs, shRNAs, transfections, adenovirus production, and chemical reagents

The DPII-FL-GFP construct and adenovirus generation using Gateway recombination was previously described whereby DPII-GFP was cloned into the pAd CMV/V5-DEST vector [17]. The DPII-GFP Carvajal construct was created by generating a point mutation in the DPII-FL-GFP construct to match the reported mutation observed in the Carvajal patient reported in Norgett et al. [31] (Epoch Life Science). The EmGFP BLOCK-iT PolIII miR RNAi Expression Vector kit (Thermo Fisher Scientific) was used to design and generate control and rat DP-specific oligonucleotides which were cloned into the pAd CMV/V5-DEST vector using Gateway recombination (target sequences: 5′-AAACCGGAAACATCATCTCTT-3′ and 5′-TGGTAATAGTTGACCCAGAAA-3′). Rat Ect2 specific oligonucleotide adenovirus was created in the same manner (5’-AGTAGGAGATGGTAACACACT-3’ and 5’-TGAAGTGTCTGCCAAGCTAGT-3’). Adenovirus was generated using the ViraPower Adenoviral Expression System (Invitrogen). Transient transfections were carried out using DharmaFECT (GE Life Sciences) to introduce siRNA oligos (5’-UCAAAGUCCUGGAGCAAGA-3’, 5’-GCAUCCAGCUUCAGACAAA-3’, 5’-ACACCAAGAUCGCUCAGAA-3’, 5’-GUGCAGAACUUGGUAAACA-3’; Thermo Fisher siGENOME) to silence DP in HL-1 cells. Cells were plated at 60–70% confluence and incubated with DNA and DharmaFECT reagent premixed in serum-free media. Fresh medium was added 24 h after transfection, and samples were lysed or fixed 48-72 h after transfection. The PKC activator Phorbol 12-myristate 13-acetate (PMA) was diluted in DMSO and used at a concentration of 15 nM (Abcam), the PKC inhibitor Bisindolylmaleimide (BIM) was diluted in DMSO and used at a concentration of 12.5 µM (Millipore Sigma).

### Antibodies

Primary antibodies utilized in this study are as follows: NW6 rabbit anti-DP directed against the C-terminal domain of DP [60], DP2.15 mouse antibody directed against DP (ThermoFisher), 1407 chicken anti-Pg (Aves Laboratories, mouse anti-N-Cadherin (Invitrogen), sheep anti-N-Cadherin (R&D Systems), mouse anti-α-Catenin, goat anti-α-Catenin (LS Bio), mouse anti-Rho-GTP, rabbit anti-Ect2 (sc-1005; Santa Cruz Biotechnologies), mouse anti-Ect2 (LS Bio), rabbit anti-GAPDH (Sigma-Aldrich), rabbit anti-phospho-PKC Substrate Motif (Cell Signaling Technology), JL-8 mouse anti-GFP (living Colors Clontech). Peroxidase-conjugated anti-mouse, -rabbit, and –chicken secondary antibodies were used for Western blot analysis (Kirkegaard & Perry Laboratories, Inc.). Alexa Fluor 488/568/647-conjugated goat or donkey anti-mouse, -rabbit, and –chicken -goat secondary antibodies were used for immunofluorescence assays (Invitrogen).

### Western Blotting and Coimmunoprecipitation Assays

For analysis of protein expression levels, cells were washed once with PBS and subsequently lysed in urea sample buffer (8 M deionized urea, 1% sodium dodecyl sulfate, 10% glycerol, 60 mM Tris, pH 6.8, and 5% β-mercaptoethanol). Samples were loaded based on total protein and samples were run on 7.5-12% SDS PAGE gels. Gels were transferred to nitrocellulose or polyvinylidene fluoride (PVDF) membranes and probed with primary and secondary antibodies directed against proteins of interest.

For coimmunoprecipitation assays cells were washed twice with PBS and lysed with modified RIPA buffer (500 mM NaCl, 50 mM Tris, pH 7.6, 10 mM MgCl_2_, 1% Triton X-100, 0.1% SDS, and 0.5% deoxycholate) supplemented with complete protease inhibitor cocktail (Millipore Sigma, Roche). Samples were then incubated with antibody against the protein of interest to be pulled down, rotated overnight at 4°C. Subsequently, samples were incubated with Protein A/G PLUS Agarose beads (Santa Cruz Biotechnology) for 30 minutes at 4°C or samples were incubated with RhoA G17A beads for active RhoA pulldowns. Samples were washed and eluted in Laemmli buffer with 5% β-mercaptoethanol, followed by SDS PAGE and Western blot analysis.

### GFP Trap

Virally infected NHEKs were cultured in 1.2 mM CaCl_2_ containing media 48h and lysed in NP40 buffer (10 mM Tris–HCl pH8, 100 mM NaCl, 0.2% NP40, 10% glycerol, with protease inhibitor tablet) and the GFP trap experiment was performed following manufacturer’s instructions (Chromotek).

### Proximity Ligation Assay (PLA)

Reagents for PLA were purchased from Sigma-Aldrich. For the proximity ligation assay (PLA) protocol, the immunofluorescence protocol described for 2D and tissue samples was followed through the addition of primary antibody. Following primary antibody incubation, the PLA protocol for 2D and tissue samples was performed according to manufacturer’s instructions. Additional information on troubleshooting, image acquisition, and analysis is detailed in Hegazy et al. [61]. Samples were mounted with PLA mounting medium containing DAPI and ImageJ software was used to quantify the percent area of PLA signal per field.

### Immunofluorescence

Cells grown on glass coverslips were washed once with PBS then fixed using 4% paraformaldehyde or ice-cold methanol. Cells were then permeabilized with 0.2% Triton X-100 in PBS. For Rho-GTP staining fixation was performed with ice-cold 10% trichloroacetic acid (TCA) in water for 15 min followed by a 20-min extraction in ice cold 0.2% Triton X-100. Primary antibodies were added to coverslips and incubated at 37°C for 1 h followed by multiple washes in PBS. Alexa fluor secondary antibodies (Thermo Fisher) were then added and incubated for 30 min, followed by PBS washes and mounting of coverslips in ProLong Gold (Thermo Fisher). Images were acquired using an AxioVison Z1 system (Carl Zeiss) with Apotome slide module, an AxioCam MRm digital camera, and either a 20x (0.8 NA Plan-Apochromat), 40x (1.4 NA, Plan-Apochromat, oil objective) or 100x (1.4 NA, Plan-Apochromat). 3D-SIM was performed using an N-SIM Super Resolution Microscope (Nikon N-SIM, Nikon, Tokyo, Japan) using an oil immersion objective lens (CFI SR Apochromat 100x, 1.49 NA, Nikon). 9 optical sections were taken at a step size of 0.2µm. Nikon Elements Advanced Research with an N-SIM module was used to reconstruct the structured illumination images. Illumination contrast modification, high-resolution noise suppression, and out-of-focus blur suppression were set with values for image reconstruction as in the Table below.

|  | 488 | 561 | 640 | 405 |
| --- | --- | --- | --- | --- |
| Illumination Contrast Modification | 0.51 | 1.02 | 0.50 | 1.00 |
| High Frequency Noise Suppression | 1.13 | 0.56 | 1.00 | 0.56 |
| Out-of-focus Blur Suppression | 0.18 | 0.17 | 0.1 | 0.11 |

### Immunohistochemistry

Tissue samples were fixed with 10% neutral-buffered formalin, embedded in paraffin and cut into 4-5 µm sections. Paraffin embedded sections were baked at 60°C overnight and deparaffinized using xylene. Samples were run through a series of ethanol and PBS dips and slides were permeabiized in 0.5% Triton X-100 in PBS. Antigen retrieval for paraffin-embedded sections was performed by heating samples to 95°C in 0.01 M citrate buffer. Sections were blocked in blocking buffer (1% Bovine serum albumin, 2% normal goat serum in PBS) and incubated with primary and secondary antibodies at 37°C. After mounting using ProLong Gold (Thermo Fisher) sections were visualized as described above using an AxioVison Z1 system (Carl Zeiss) with Apotome slide module, an AxioCam MRm digital camera.

### Statistics

All quantifications are presented as the mean ± SEM unless otherwise stated. One way ANOVA followed by Tukey’s post-hoc analysis or Dunnett’s multicomparison test was used for multiple comparisons. For data comparing two conditions, statistical differences were analyzed with a two tailed Student’s *t* test. P<0.05 was considered significant. For statistical analysis presented as fold change (bar graphs), the mean of data from control samples was assigned a value of 1 and all other samples were calculated relative to the control. All densitometry quantification was normalized to respective loading controls.

## Acknowledgements

The authors would like to acknowledge Jeffrey Saffitz for contributing patient sections. The work was supported by the following grants: NIH R01 AR041836, R01 AR043380, and R01 CA228196 and the J.L. Mayberry Endowment to KJG; American Heart Association Fellowships to C.K and H.Z; NIH/NIAMS F32AR081677 to A.P, NIH R01HL142251, NIHR01 HL162369 and LEXEO Therapeutics Inc to FS. F.S. was a co-founder of Stelios Therapeutics Inc. (acquired by LEXEO Therapeutics Inc.) and is a co-founder and shareholder of Papillon Therapeutics Inc and MyoTherapeutix Inc as well as is a consultant and shareholder of LEXEO Therapeutics Inc.

## Figure Legends

**Supplementary Figure 1: Ect2 co-localizes with DP in control tissue from mouse and human hearts**. (A) Isolated rat cardiac myocyte (NRVCM) SIM images with maximum image projections of single fluorophores and orthogonal planes illustrating the colocalization of Ect2 with junction proteins. Images are associated with the merged images in Figure 1A. Scale bar = 5μm. Sections from control mouse hearts (B) and control human hearts (C) were prepared for immunofluorescence and stained with antibodies directed against DP or Ect2 and imaged using the AxioVison Z1 system with Apotome slide module. Scale bar = 20 μm. (D) Immunoprecipitation of DP was performed on isolated primary normal human keratinocytes and lysates were probed for DP and Ect2. MW lane = molecular weight ladder.

**Supplementary Figure 2: Active RhoA at IDs is decreased in *Dsp* cKO animals**. Quantification of *Dsp* cKO mouse hearts stained for DP and active Rho (Rho-GTP) shown in Figure 4B and B’. Data cloud represents intensity measurements of every ROI of a given condition; horizontal bars represent median ± standard deviation.

**Supplementary Figure 3: Ect2 junctional localization is disrupted in keratinocytes from Carvajal patients**. (A) JD-1 keratinocytes isolated from a Carvajal patient were fixed and double-stained for Ect2 and DP. Scale bar = 20μm. (B) Quantification of fluorescence intensity showed that JD-1 cells had significantly less Ect2 co-localizing with DP than Ctl adult keratinocytes (IHEK, p=0.05). (C) GFP Ctl, DPII and DPII Carvajal (CJ) GFP constructs were isolated in a GFP trap pull down and blotted back for GFP and Ect2.

