## Supplemental Figures for "Association of RhoGEF Ect2 with Desmoplakin Supports RhoA Activity at Intercellular Junctions: Implications for Carvajal Disease"

#### Supplementary Materials

[#]Northwestern Feinberg School of Medicine, Department of Pathology, 303 E. Chicago Ave., Chicago, IL. 60611

[%] Blizard Institute, Faculty of Medicine and Dentistry, Queen Mary University of London, London E1 4NS

[&] University of California-San Diego, School of Medicine, Department of Medicine, La Jolla CA 92093

*These authors contributed equally

**Corresponding authors

#### Supplementary Figure 1

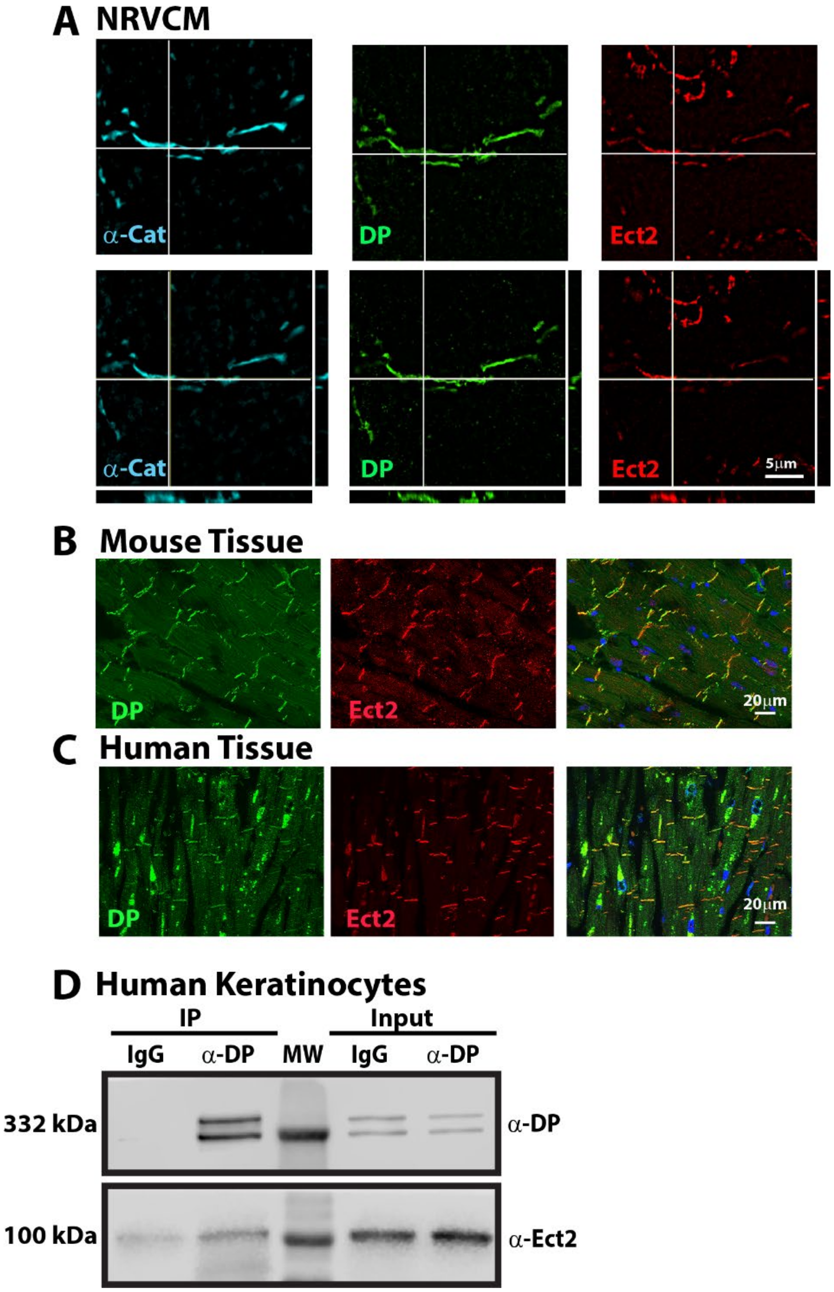


**Supplementary Figure 1: Ect2 co-localizes with DP in control tissue from mouse and human hearts.** (A) Isolated rat cardiac myocyte (NRVCM) SIM images with maximum image projections of single fluorophores and orthogonal planes illustrating the colocalization of Ect2 with junction proteins. Images are associated with the merged images in Figure 1A. Scale bar = 5μm. Sections from control mouse hearts (B) and control human hearts (C) were prepared for immunofluorescence and stained with antibodies directed against DP or Ect2. and imaged using the AxioVison Z1 system with Apotome slide module. Scale bar = 20 μm. (D) Immunoprecipitation of DP was performed on isolated primary normal human keratinocytes and lysates were probed for DP and Ect2. MW lane = molecular weight ladder.

### Supplementary Figure 2

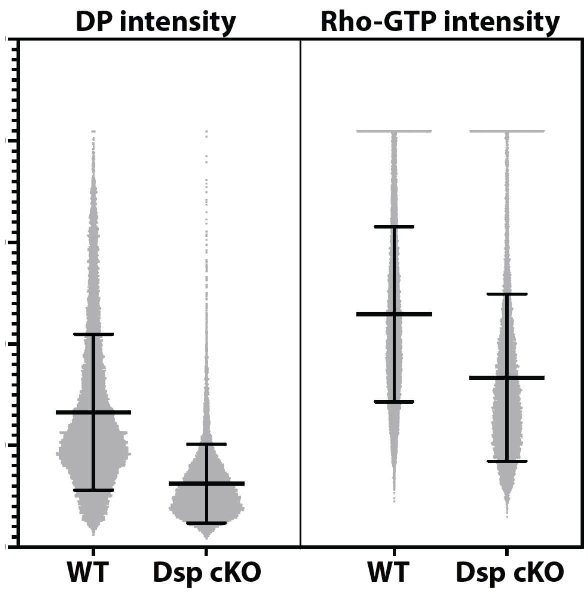


**Supplementary Figure 2: Active RhoA at IDs is decreased in *Dsp* cKO animals.** Quantification of *Dsp* cKO mouse hearts stained for DP and active Rho (Rho-GTP) shown in Figure 4B and B'. Data cloud represents intensity measurements of every ROI of a given condition; horizontal bars represent median ± standard deviation.

#### Supplementary Figure 3

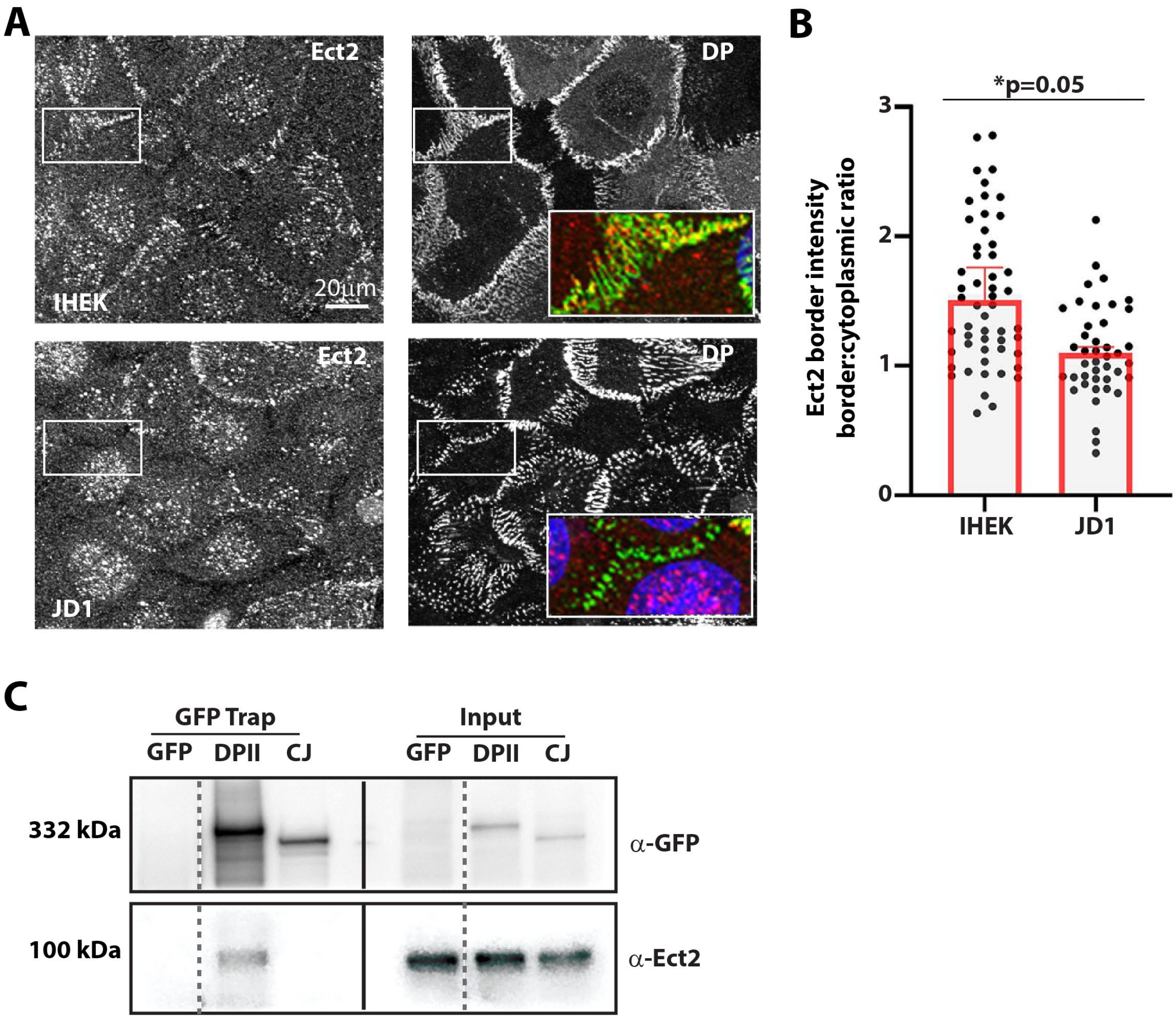


**Supplementary Figure 3: Ect2 junctional localization is disrupted in keratinocytes from Carvajal patients.** (A) JD-1 keratinocytes isolated from a Carvajal patient were fixed and double-stained for Ect2 and DP. Scale bar = 20µm. (B) Quantification of fluorescence intensity showed that JD-1 cells had significantly less Ect2 co-localizing with DP than Ctl adult keratinocytes (IHEK, p=0.05). (C) GFP Ctl, DPII and DPII Carvajal (CJ) GFP constructs were isolated in a GFP trap pull down and blotted back for GFP and Ect2.
